# Extended frontal networks for visual and auditory working memory

**DOI:** 10.1101/2021.04.16.439914

**Authors:** Abigail L. Noyce, Ray W. Lefco, James A. Brissenden, Sean M. Tobyne, Barbara G. Shinn-Cunningham, David C. Somers

## Abstract

Working memory (WM) supports the persistent representation of transient sensory information. Visual and auditory stimuli place different demands on WM and recruit different brain networks. Separate auditory- and visual-biased WM networks extend into the frontal lobes, but several challenges confront attempts to parcellate human frontal cortex, including fine-grained organization and between-subject variability. Here, we use differential intrinsic functional connectivity from two visual-biased and two auditory-biased frontal structures to identify additional candidate sensory-biased regions in frontal cortex. We then examine direct contrasts of task fMRI during visual vs. auditory 2-back WM to validate those candidate regions. Three visual-biased and five auditory-biased regions are robustly activated bilaterally in the frontal lobes of individual subjects (N=14, 7 women). These regions exhibit a sensory preference during passive exposure to task stimuli, and that preference is stronger during WM. Hierarchical clustering analysis of intrinsic connectivity among novel and previously identified bilateral sensory-biased regions confirms that they functionally segregate into visual and auditory networks, even though the networks are anatomically interdigitated. We also observe that the fronto-temporal auditory WM network is highly selective and exhibits strong functional connectivity to structures serving non-WM functions, while the fronto-parietal visual WM network hierarchically merges into the multiple-demand cognitive system.

---

Sensory working memory (WM) recruits a network of brain structures that spans frontal, and posterior cortical areas as well as sub-cortical regions (Arnott et al., 2005; Brissenden et al., 2018; Christophel et al., 2012; Crottaz-Herbette et al., 2004; Harrison & Tong, 2009; Huang et al., 2013; Jerde et al., 2012; Kastner et al., 2007; Lewis-Peacock et al., 2015; Michalka et al., 2015; Owen et al., 2005; Postle et al., 2000). Each sensory modality receives information from a unique set of receptors that is then processed by modality-specific subcortical and primary cortical regions, and each sensory modality possesses distinct strengths and weaknesses in the resolution and fidelity with which information can be encoded (Alais & Burr, 2004; Noyce et al., 2016; Welch & Warren, 1980). One leading model of WM has proposed multiple WM components, including specialized visuospatial and auditory/phonological stores (Baddeley, 2010; Baddely & Hitch, 1974). Although frontal lobe WM mechanisms have long been viewed as agnostic to sensory modality (Assem et al., 2020; Duncan, 2010; Duncan & Owen, 2000; Fedorenko et al., 2013; Postle et al., 2000; Tamber-Rosenau et al., 2013), a growing body of research reveals substantial influences of sensory modality on the functional organization of WM structures throughout the brain, even in frontal cortex (Hagler & Sereno, 2006; Kastner et al., 2007; Kumar et al., 2016; Mayer et al., 2016; Michalka et al., 2015; Noyce et al., 2017; Romanski, 2007; Romanski & Goldman-Rakic, 2002). Work from our laboratory has previously identified both sensory specialized areas for WM (Michalka et al., 2015; Noyce et al., 2017; Tobyne et al., 2017) as well as areas that seem to be recruited independent of sensory modality (Noyce et al., 2017). Although these regions are relatively modest in size, their characteristic pattern of organization is repeated across individual subject cortical hemispheres. Moreover, these sensory-biased frontal lobe regions form specific functional connectivity networks with traditionally identified sensory-specific regions in parietal and temporal cortex.

In prior work examining sensory-biased attention and WM structures in frontal lobes, we reported two visual-biased and two auditory-biased regions in nearly all individual subject hemispheres (Michalka et al., 2015; Noyce et al., 2017). We subsequently leveraged our small in-lab dataset to examine sensory-biased functional connectivity networks (Tobyne et al., 2017) by using resting-state fMRI data from 469 subjects of the Human Connectome Project (Glasser et al., 2016). This analysis examined differential functional connectivity from parieto-occipital visual and temporal auditory attention regions to frontal cortex. The results not only confirmed that the previously identified sensory-biased frontal regions reside within sensory-specific functional networks, but also suggested that these sensory-specific networks might extend more anteriorly on the lateral surface of frontal cortex and to the medial surface of frontal cortex (Tobyne et al., 2017). However, these analyses were performed with group data, and without the benefit of sensory-specific WM task activation in individual subjects. Here, we investigate more fully whether and how these sensory-biased attention networks might extend within frontal cortex.

We are motivated to examine this question for three reasons. First, our previous results focused on caudolateral frontal cortex, ignoring evidence of specialization in frontal opercular regions as well as on the medial surface. Second, we know from both our and other’s work that small regions whose exact anatomical location varies across subjects can be difficult to identify and characterize (Braga & Buckner, 2017; Mueller et al., 2013; Tobyne et al., 2018). Third, we previously observed variability within regions that we characterized as “multiple demand,” and now seek to more closely examine the functional properties of these different regions.

We used differential functional connectivity (Lefco et al., 2020; Tobyne et al., 2017) from sensory-biased frontal seed regions to identify thirteen candidate sensory-biased regions bilaterally throughout the frontal lobe. After assessing their reliability in individual subjects’ task activation, we drew subject-specific labels for eight bilateral regions. These were further characterized in terms of (1) their sensory and WM recruitment; (2) their consistency of location across individual subjects; and (3) their network organization. We found that three bilateral visual-biased WM regions and five bilateral auditory-biased WM regions in frontal cortex occur with high consistency in individual subjects. These regions exhibit sensory-biased activation both during passive exposure to visual and auditory stimuli, and during 2-back WM.

## Methods

All procedures were approved by the Institutional Review Board of Boston University.

### Overview

After collecting task (visual and auditory, WM and sensorimotor control) and resting fMRI data, we drew subject-specific labels for two bilateral visual-biased and two bilateral auditory-biased regions (Figure 1A). These served as seeds for the differential connectivity analysis. From the group-level differential connectivity results, we defined candidate sensory-biased regions, whose task activation was then assessed in individual subjects. Eight regions met our criteria for investigation via individual subject labels (Figure 1B); for those, we report their typical location and size, their degree of sensory and WM-specific recruitment, and their network organization structure. Five candidate regions were not reliable in individual subjects (Figure 1C); we report sensory-specific WM recruitment in each of these rejected candidates.

**Figure 1.**
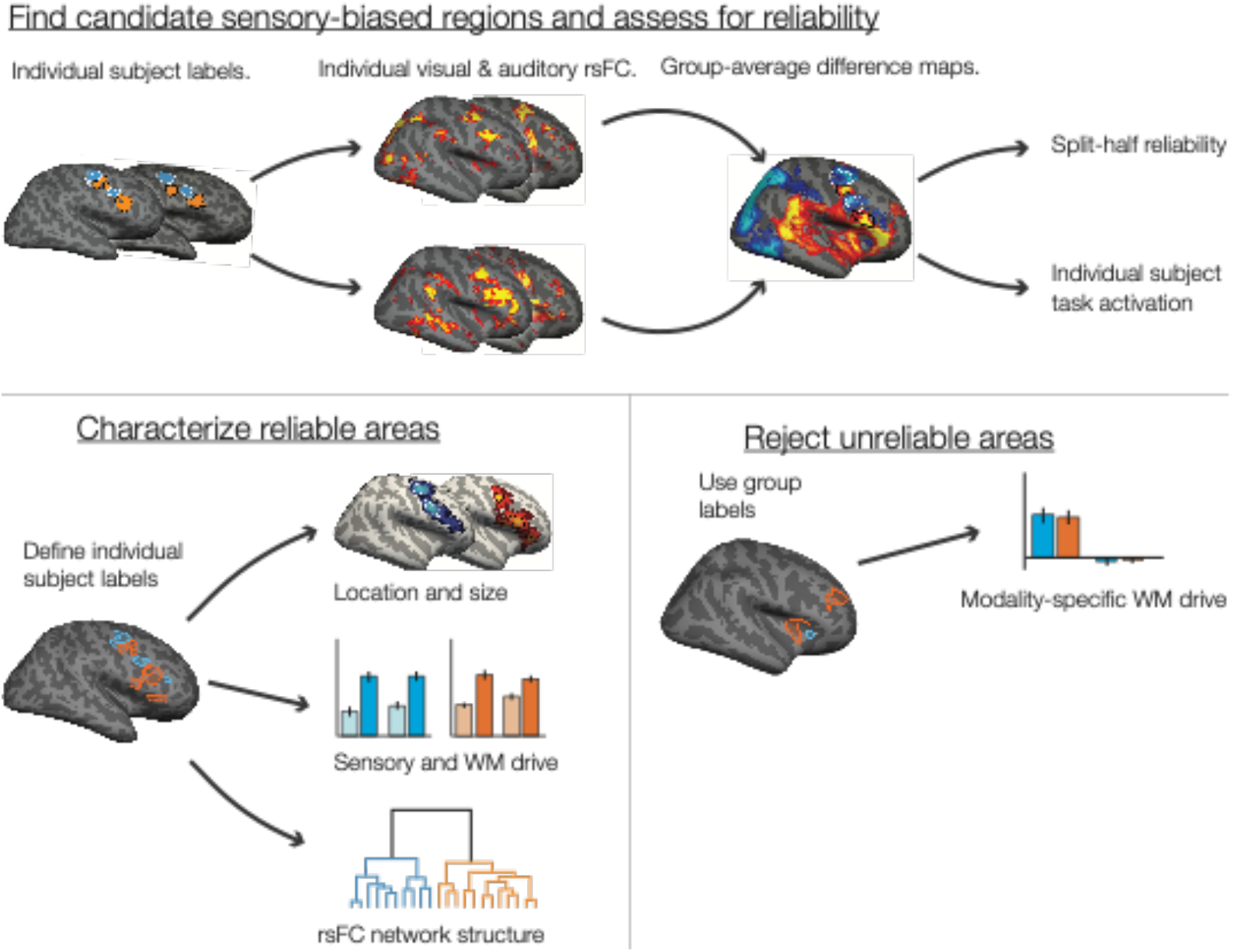
Overview of analytical steps. (A) We began with previously-identified sensory-biased regions in caudolateral frontal cortex, along the precentral sulcus and inferior frontal sulcus (Michalka et al., 2015; Noyce et al., 2017). Within each individual subject, that subject’s auditory- and visual-biased regions were used as seeds to compute seed-to-whole-hemisphere rsFC. We thresholded (see Methods) and z-transformed the resulting connectivity maps before taking the difference between auditory-biased and visual-biased connectivity. The resulting differential connectivity maps were used to identify candidate sensory-biased regions (Figure 3A). We assessed each candidate region using split-half reliability of task activation (Figure 3B) and hand-scoring the region’s appearance in individual subjects (Table S1). (B) Regions that exhibited consistent sensory-biased WM recruitment both within and across subjects were denoted as reliable regions; we drew subject-specific labels for each region in each subject for analysis (Figure 4). We reported the mean location and size (Figure 4 and Table 1), the degree of WM-specific recruitment (Figure 5) and used hierarchical clustering of seed-to-seed functional connectivity to investigate the structure of sensory-biased networks (Figure 6). (C) Regions that did not exhibit consistent sensory-biased recruitment were investigated using the candidate search space labels; we again reported the degree of WM-specific recruitment in each (Figure 7).

### Subjects

Sixteen members of the Boston University community (ages 24–35; nine men and seven women) participated in this study. All experiments were approved by the Institutional Review Board of Boston University. All subjects gave written informed consent to participate and were paid for their time. Two authors (ALN & JAB) participated as subjects. One subject was excluded from analysis due to technical difficulties with the auditory stimulus presentation; another participated in task scans but not resting-state scans and is included only in task activation analyses.

### Experimental task

Subjects performed a 2-back WM task for visual (photographs of faces) and auditory (recordings of animal sounds) stimuli, in separate blocks (Figure 2). Each block contained 32 stimuli and lasted 40 seconds; onsets of successive stimuli were 1.25 s apart. Visual stimuli were each presented for 1 s; auditory stimuli ranged from 300-600 ms in duration. Images were presented at 300×300 pixels, spanning approximately 6.4° visual angle, using a liquid crystal display projector illuminating a rear-projection screen within the scanner bore. Auditory stimuli were natural recordings of cat and dog vocalizations. Stimuli were presented diotically. The audio presentation system (Sensimetrics, http://www.sens.com) included an audio amplifier, S14 transformer, and MR-compatible in-ear earphones. At the beginning of each block, subjects were cued to perform either the visual or the auditory 2-back task, or to perform a sensorimotor control condition. Each run comprised 8 32-second blocks: 2 auditory 2-back, 2 visual 2-back, 2 auditory sensorimotor control, and 2 visual sensorimotor control. Eight seconds of fixation was recorded at the beginning, midpoint, and end of each run. Block order was counterbalanced across runs; run order was counterbalanced across subjects. During 2-back blocks, participants were instructed to decide whether each stimulus was an exact repeat of the stimulus two prior, and to make either a “2-back repeat” or “new” response via button press. Sensorimotor control blocks consisted of the same stimuli and timing, but no 2-back repeats were included, and participants were instructed to make a random button press to each stimulus. Responses were collected using an MR-compatible button box. All stimulus presentation and task control was managed by custom software using Matlab PsychToolbox (Brainard, 1997; Cornelissen et al., 2002; Kleiner et al., 2007).

**Figure 2.**
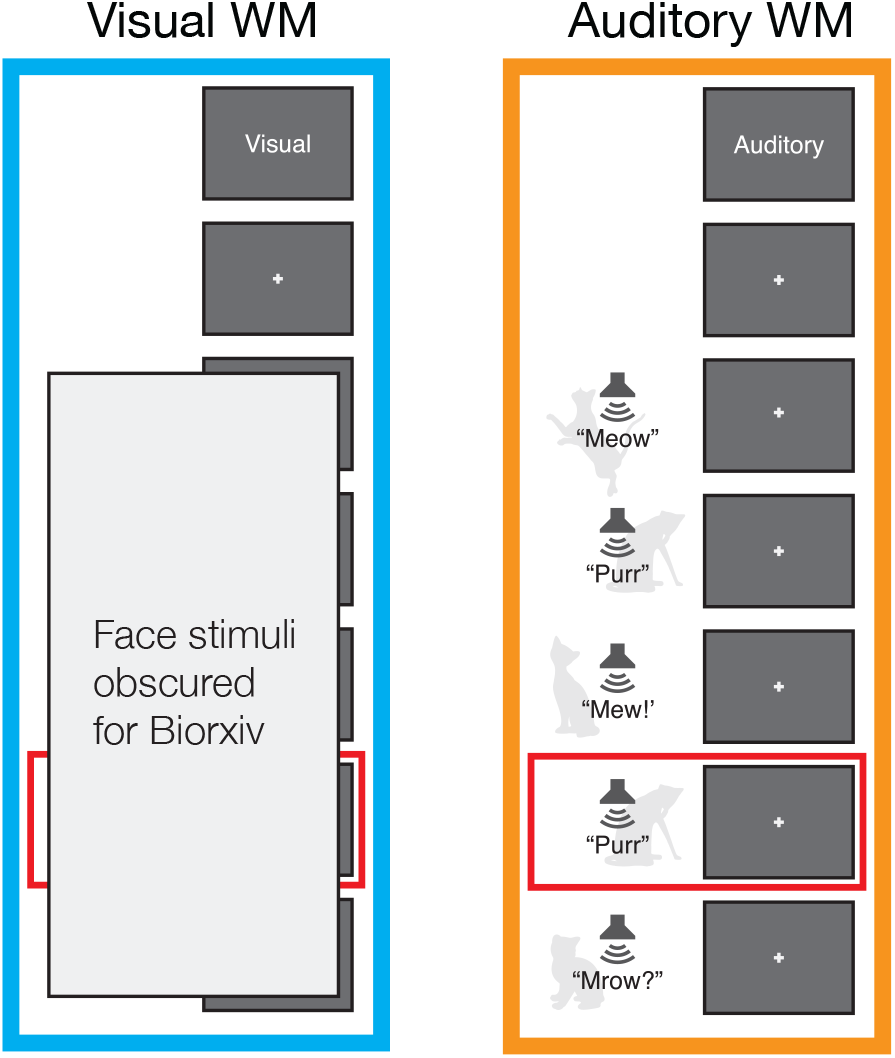
Diagram of the two-back WM task used to identify and characterize sensory-biased and sensory-independent WM structures. Subjects observed a stream of visual or auditory stimuli and reported 2-back repeats via button-press. Sensorimotor control blocks consisted of the same stimuli, without repeats included. (A) The visual stimuli comprised back-and-white photographs of faces, drawn from a corpus of student ID photos. (B) The auditory stimuli comprised recordings of cat and dog vocalizations.

Subjects (*n* = 15) were able to perform both the visual 2-back task (90.1% correct, SD = 7.5%) and the auditory 2-back task (87.5% correct, SD = 10.9%) at a high level. There was no significant difference in accuracy between the two tasks (*t*(14) = 1.463, *p* = 0.17).

### Eye tracking

All subjects were trained to hold fixation at a central point. Eye movements during task performance were recorded using an Eyelink 1000 MR- compatible eye tracker (SR Research) sampling at 500 Hz. Eye tracking recordings were unavailable for four subjects due to technical problems; eye gaze in these subjects was monitored via camera to confirm acceptable fixation performance. We operationalized fixation as eye gaze remaining within 1.5° visual angle of the central fixation point. Eye position data were smoothed through an 80-Hz low-pass first-order Butterworth filter, after which excursions from fixation were counted within each block. To rule out differences in eye movements as a potential confound, we compared the frequency with which subjects broke fixation between visual and auditory 2-back blocks, between visual and auditory sensorimotor control blocks, and between each sensory modality’s 2-back and sensorimotor control blocks. There were no significant differences in any of these comparisons (paired t-tests, all *p* > .35).

### MR Imaging and Analysis

#### MR scanning

All scans were performed on a Siemens TIM Trio 3T MR imager, with a 32-channel matrix head coil. Each subject participated in multiple MRI scanning sessions, including collection of high-resolution structural images, task-based functional MRI, and resting-state functional MRI. High-resolution (1.0 × 1.0 × 1.3 mm) magnetization-prepared rapid gradient echo (MP-RAGE) T1-weighted structural MR images were collected for each subject. Functional T2*-weighted gradient-echo echo-planar images were collected using a slice-accelerated EPI sequence that permits simultaneous multi-slice acquisition via the blipped-CAIPI technique (Setsompop et al., 2012). Sixty-nine slices (0% skip; TE 30 ms; TR 2000 ms; 2.0 × 2.0 × 2.0 mm) were collected with a slice acceleration factor of 3. Partial Fourier acquisition (6/8) was used to keep TE in a range with acceptable signal to noise ratios. We collected 8 runs of task-based functional data per subject and we collected 2 to 3 runs (360 seconds per run) of eyes-open resting state functional data per subject. MRI data collection was performed at the Harvard University Center for Brain Science neuroimaging facility. The structural and task data reported here were also included in a previous report (Noyce et al., 2017).

#### Structural processing

The cortical surface of each hemisphere was reconstructed from the MP- RAGE structural images using FreeSurfer software (http://surfer.nmr.mgh.harvard.edu, Version 5.3.0, details in Fischl, Sereno, Tootell, & Dale, 1999; Dale, Fischl, & Sereno, 1999; Fischl, Sereno, & Dale, 1999; Fischl, Liu, & Dale, 2001; Fischl, 2004). Each cortical reconstruction was manually checked for accuracy.

#### Functional preprocessing and GLM

Functional data were analyzed using FreeSurfer’s FS-FAST analysis tools (version 5.3.0). Analyses were performed on subject-specific anatomy unless noted otherwise. All task and resting state data were registered to individual subject anatomy using the middle time point of each run. Data were slice-time-corrected, motion corrected by run, intensity normalized, resampled onto the individual’s cortical surface (voxels to vertices) and spatially smoothed on the surface with a 3mm FWHM Gaussian kernel. Task data were also resampled to the FreeSurfer *fsaverage* surface for group analysis.

To analyze task data, we used standard functions from the FS-FAST pipeline to fit a general linear model (GLM) to each cortical vertex. The regressors of the GLM matched the time course of the experimental conditions. The time points of the cue period were excluded by assigning them to a regressor of no interest. The canonical hemodynamic response function was modeled by a gamma function (δ = 2.25 s, τ = 1.25); this hemodynamic response function was convolved with the regressors before fitting.

Resting-state data (N = 14) were collected using the same T2*-weighted sequences that were used to collect task fMRI. For each subject, we collected 2 to 3 runs (720–1080 s, 360-540 timepoints) of eyes-open resting state, during which subjects maintained fixation at the center of the display. Data were preprocessed similarly to the task data: slice-time corrected, motion-corrected by run, intensity normalized, resampled onto the individual’s cortical surface, and spatially smoothed on the surface with a 3mm FWHM Gaussian kernel. Multiple resting-state acquisitions for each subject were temporally demeaned and concatenated to create a single timeseries. In order to attenuate artifacts that could induce spurious correlations, resting-state data were further preprocessed using custom scripts in MATLAB. The following preprocessing steps were performed: linear interpolation across high-motion time-points (> 0.5 mm framewise displacement (FD); (Carp, 2013; Power et al., 2012)), application of a fourth-order Butterworth temporal bandpass filter to extract frequencies between 0.009 and 0.08 Hz, mean ‘grayordinate’ signal regression (Burgess et al., 2016), and censoring of high-motion time-points by deletion (Power et al., 2012).

#### Differential connectivity

In order to identify candidate sensory-biased structures within the hypothesized extended networks, we computed differential functional connectivity (Lefco et al., 2020; Tobyne et al., 2017) from previously-defined visual-biased (superior and inferior precentral sulcus, sPCS and iPCS) and auditory-biased (transverse gyrus bridging precentral sulcus, tgPCS, and caudal inferior frontal sulcus/gyrus, cIFS/G) frontal seeds (Figure 1A). For each subject, the two visual-biased frontal labels from Noyce et al., (2017) were combined into one single functional-connectivity seed per hemisphere, as were the two auditory-biased frontal labels. For each seed in each individual subject hemisphere, we calculated a mean resting-state time course over all vertices within the seed. We then computed the correlation (Pearson’s r) between the two seeds’ timecourses and those of every vertex within the same cortical hemisphere.

Visual seed and auditory seed correlation maps for each subject hemisphere were resampled to the FreeSurfer *fsaverage* surface and subjected to Fisher’s *r*-to-*z* transformation, before submission to a group-level analysis. Group-average *z* maps were standardized within each cortical hemisphere by subtracting the mean *z*-value from each vertex, and then dividing by the standard deviation. Difference maps were then created by subtracting the visual-seed connectivity map from the auditory-seed connectivity map (Figure 3A).

**Figure 3.**
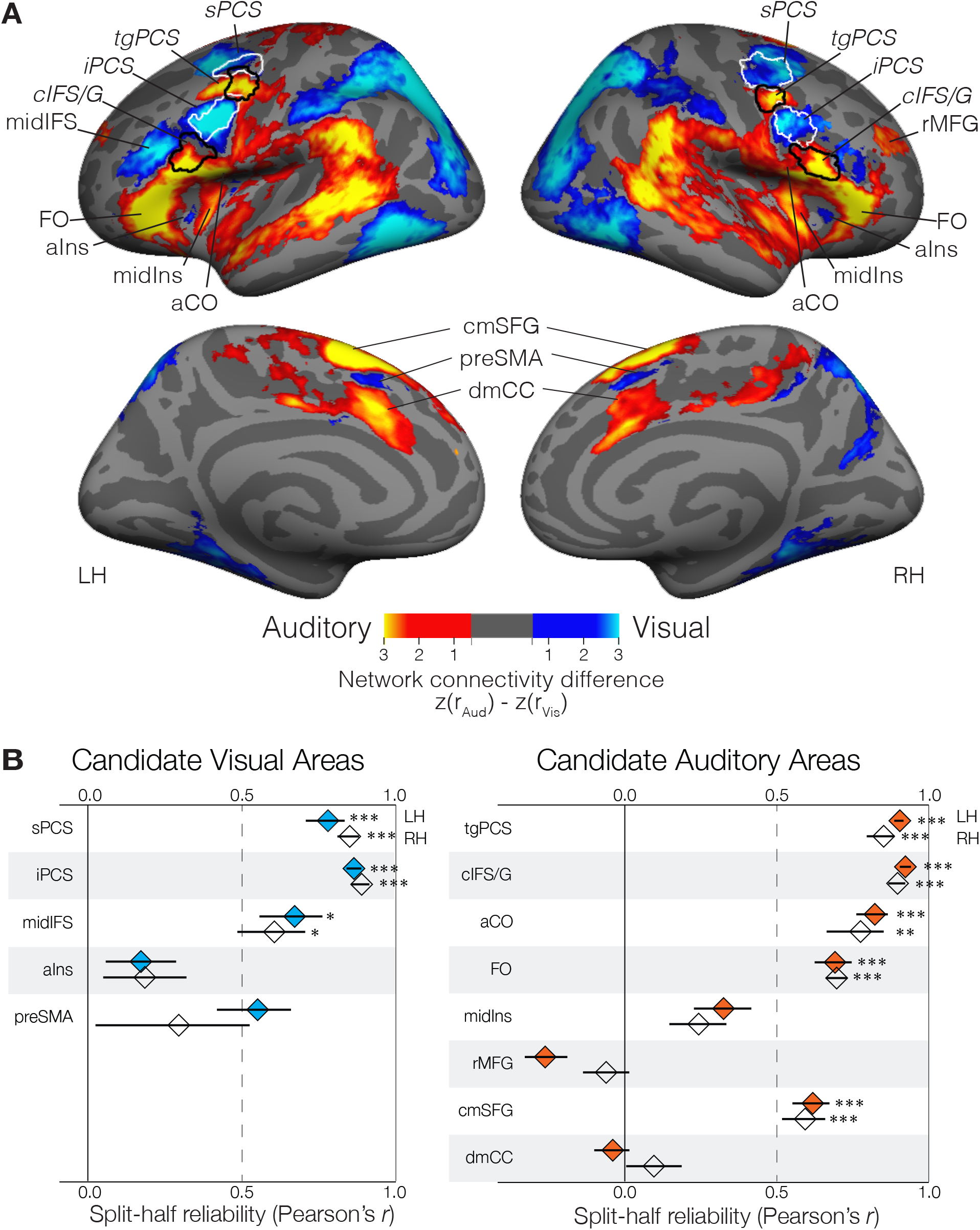
(A) Group average (N=14) differential functional connectivity to visual-biased frontal regions (seeds outlined in white; significantly greater connectivity shown in blue) and auditory-biased frontal regions (seeds outlined in back; significantly greater connectivity shown in yellow). The four bilateral seed regions were defined in individual subjects from task activation (Seed label outlines shown here are regions where individual subject labels overlap in at least 20% of subjects.) Note clear preferential connectivity between visual frontal seeds and posterior visual attention regions (parietal and occipital cortex) and between auditory frontal seeds and posterior auditory attention regions (superior temporal cortex). Within frontal cortex, we observe regions of preferential visual network connectivity to bilateral aIns and preSMA as well as to left midIFS; we observe regions of preferential auditory connectivity to bilateral aCO, FO, midIns, cmSFG, and dmCC, as well as to right rMFG. Maps are thresholded at p<.05 after correcting for multiple comparisons (Smith & Nichols, 2009). (B) Split-half reliability (Pearson’s r) of task activation (Aud WM vs. Vis WM) for each candidate visual-biased (left, blue) and auditory-biased (right, orange) region. Three bilateral visual-biased regions (sPCS, iPCS, and midIFS) and five bilateral auditory-biased regions (tgPCS, cIFS/G, aCO, FO, and cmSFG) are significantly non-zero. Left preSMA is above r = 0.5 but does not survive multiple comparisons correction. Error bars are within-subject standard error of the Fisher’s z-transformed mean correlation (Cousineau, 2005; Loftus & Masson, 1994; Morey, 2008). *p < .05, **p < .01, ***p < .001, Holm-Bonferroni corrected for 26 tests.

To control family-wise error rate, we restricted this map to those vertices that exhibited significant visual or auditory connectivity after multiple comparisons correction. We used permutation testing to generate a null distribution of threshold free cluster enhancement (TFCE)(Smith & Nichols, 2009), and compared the TFCE values of the group average visual and auditory connectivity maps to this distribution. We thresholded at p < .05, one-sided; the union of vertices that survived corrections in the visual and auditory connectivity maps was used to mask the final differential connectivity map. From this difference map, we identified seven new bilateral and two new unilateral candidate sensory-biased frontal regions, in addition to the original four bilateral seeds.

#### Assessing candidate regions

We used two methods to determine which candidate sensory-biased frontal regions (from the differential connectivity analysis) were reliable enough to characterize. First, we computed split-half reliability of task activation in each region. Group-space labels from the differential connectivity analysis (Figure S1) were projected back into individual subjects. Where a region only appeared unilaterally, the mirrored label was also created. Each individual subject’s data were divided into odd- and even-numbered runs, and a univariate first-level analysis was performed as described above. Vertices within each candidate region were masked to include only those which showed the expected direction of activation in at least one set of runs (to exclude any contribution from adjacent, opposite-biased regions). We then computed the vertex-wise correlation (Pearson’s *r*) between odd-numbered and even-numbered runs. Correlation values were Fisher’s *z*-transformed for averaging and computing standard error, and transformed back to *r* values for visualization (Figure 3B).

Second, two authors (A.L.N and D.C.S.) visually inspected task-activation maps of the univariate first-level auditory 2-back versus visual 2-back contrast (see below). We independently scored each ROI’s strength in each subject, considering its size, activation intensity, position, and compactness (Table S1). Regions were scored as strong (1), weak (0.5), or absent (0). Again, although two regions only appeared unilaterally in the corrected differential connectivity map, we assessed them in both hemispheres at this stage.

#### Task activation and labels

For each subject, we directly contrasted blocks in which the subject performed visual WM against blocks in which the subject performed auditory WM. This contrast was liberally thresholded at p < .05, uncorrected. This threshold was set to maximally capture frontal lobe vertices showing a bias for auditory or visual WM. For regions that occurred reliably in individual subjects (mean correlation above 0.5 and mean scoring above 0.7), we drew labels based on each individual subject’s activation pattern in this contrast. This resulted in a set of three bilateral visual-biased frontal regions and five bilateral auditory-biased frontal regions for further analysis. Note that sPCS, iPCS, tgPCS, and cIFS/G were previously identified as sensory-biased frontal regions (Michalka et al., 2015; Noyce et al., 2017) and were used as seeds for the differential connectivity analysis, above. We also drew large posterior visual (pVis, including parietal and occipital regions) and posterior auditory (pAud, including superior temporal gyrus and sulcus) labels for each subject, to capture canonical sensory attention structures; these labels were used in the functional connectivity analysis (see below).

Within each region, we computed mean % BOLD signal change in two contrasts: auditory sensorimotor control versus visual sensorimotor control, and auditory 2-back versus visual 2-back (Figure 1B, Figure 4A). The first contrast captures the degree of sensory drive in each region in the absence of the WM task; the difference between the two contrasts captures WM specific activation.

**Figure 4.**
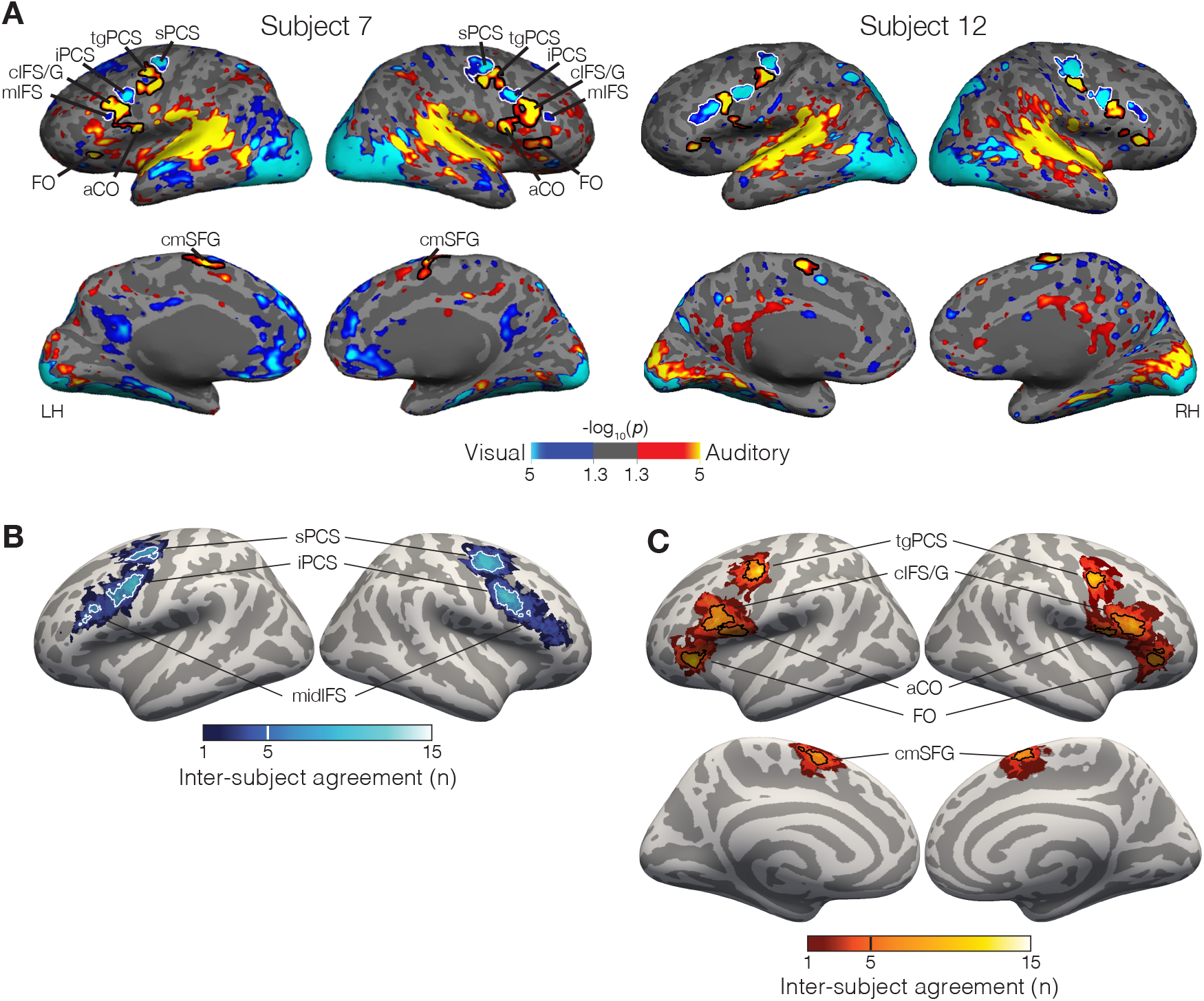
(A) Contrast of auditory 2-back (yellow) vs. visual 2-back (blue) WM task activation in two representative subjects (all subjects are shown in Figure S2). In lateral frontal cortex, we observe three visual-biased structures: superior and inferior precentral sulcus (sPCS and iPCS) and mid inferior frontal sulcus (midIFS). They are interleaved with auditory-biased structures including the transverse gyrus intersecting precentral sulcus (tgPCS), caudal inferior frontal sulcus/gyrus (cIFS/G), anterior central operculum (aCP), and frontal operculum (FO). On the medial surface we observe auditory-biased caudomedial superior frontal gyrus (cmSFG). (B) Probabilistic overlap map of visual-biased ROI labels projected into fsaverage space. Although there is great variability in the exact positioning of these regions, they overlap into three clear “hot spots”. Thick outlines are drawn around vertices that are included in 5 or more subjects (33% of the participants); for right midIFS thin outlines are drawn around vertices that are included in 3 or more subjects (20% of the participants). (C) Probabilistic overlap map of auditory-biased ROI labels projected into fsaverage space. Outlines are drawn around vertices that are included in 5 or more subjects (33% of the participants).

Probabilistic ROIs (Figure 4B,C) were constructed as in Tobyne et al., 2017, by projecting individual subject task activation (binarized at p < 0.05, uncorrected) to the fsaverage template surface via spherical registration and trilinear interpolation (Fischl, Sereno, & Dale, 1999). For each surface vertex, we computed the proportion of subjects for whom that vertex was included in a given region of interest.

#### Network clustering analysis

To compute seed-to-seed or seed-to-whole-brain connectivity among the ROIs, we took the mean resting-state timecourse across all vertices in each region. Pairwise correlations between timecourses measured connectivity between seeds, or between a seed and each surface vertex.

For each ROI, its connectivity to each other seed gave a connectivity profile across the network; these connectivity profiles were averaged across subjects and then fed into a hierarchical clustering analysis to characterize the network structure (Brissenden et al., 2018; Dosenbach et al., 2007; Michalka et al., 2015; Tobyne et al., 2017). We computed pairwise Euclidean distance between each region’s connectivity profile. We then applied Ward’s linkage algorithm to these distances, which forms each new cluster by merging the two clusters that lead to the minimum possible increase in the total sum of squares of the node to centroid distances. Reliability of the clusters was assessed via a bootstrap approach: on each of 10,000 iterations, fourteen datasets were sampled (with replacement) and the hierarchical clustering analysis was performed. This yielded a bootstrapped distribution of network structures. For each subtree in the original structure, we counted how frequently it appeared in the bootstrapped distribution.

## Results

### Candidate sensory-biased frontal structures identified from differential functional connectivity

We began by examining functional connectivity within frontal cortex in order to identify ‘candidate’ regions that are likely members of either visual-biased or auditory-biased WM networks. Previously, we examined functional connectivity of frontal cortex to visual-biased parieto-occipital cortex and to auditory-biased temporal cortex (Tobyne et al., 2017). Many regions in frontal cortex exhibited preferential connectivity to visual or auditory posterior cortex. That analysis, performed on publicly available HCP data, lacked the task data necessary to validate the candidate network regions (Tobyne et al., 2017).

Here, we examined differential functional connectivity within frontal cortex by contrasting connectivity from visual-biased frontal regions (sPCS and iPCS combined as a single seed; (Michalka et al., 2015; Noyce et al., 2017)) against connectivity from previously identified auditory-biased frontal regions (tgPCS and cIFS/G combined as a single seed; (Michalka et al., 2015; Noyce et al., 2017)). We constructed connectivity maps for each subject hemisphere from each seed; maps were combined across subjects (using threshold free cluster enhancement and non-parametric randomization tests to create significance masks; (Smith & Nichols, 2009)) and the two sensory-biased connectivity maps were subtracted from each other to yield a differential connectivity map (Figure 3A). The differential connectivity map shows several expected patterns. First, this contrast identifies preferential connectivity between visual-biased frontal regions and posterior parietal and occipital areas, and between auditory-biased frontal regions and posterior areas in superior temporal lobe, similar to the results of Michalka et al. (2015) and Tobyne et al. (2017). Second, this contrast recapitulates the seed regions that went into the connectivity analysis (visual-biased sPCS and iPCS; auditory-biased tgPCS and cIFS/G).

More interestingly, this contrast suggests extended sensory-biased structures within frontal, insular, and cingulate cortex. The visual-biased frontal seeds (sPCS and iPCS) are preferentially connected to bilateral regions in anterior insula (aIns) and pre-supplementary motor area (preSMA), as well as to left middle inferior frontal sulcus (midIFS). The auditory-biased frontal seeds (tgPCS and cIFS/G) are preferentially connected to bilateral regions in anterior central operculum (aCO), frontal operculum (FO), middle insula (midIns), caudomedial superior frontal gyrus (cmSFG), and dorsal middle cingulate cortex (dmCC), as well as to right rostral middle frontal gyrus (rMFG). Therefore, the differential functional connectivity analysis yields three novel (and two previously identified) frontal regions as candidates for the visual-biased network, along with six novel (and two previously identified) frontal candidates for the auditory-biased network.

### Defining sensory-biased frontal structures

To examine whether these candidate sensory-biased regions occurred reliably, we first tested whether they exhibited consistent patterns of task activation in the expected direction. Labels from the group-average differential connectivity map (Figure 3A, S1) were projected into individual subjects’ anatomical space (including mirrored labels for candidate regions that occurred unilaterally) and used to test split-half (odd- and even-numbered runs) reliability. For each half of the task runs, we computed auditory 2-back vs. visual 2-back contrasts, and computed the vertex-wise correlation of task activation across the ROI, between halves (Figure 3B). Bilaterally, candidate visual regions sPCS, iPCS, and midIFS exhibited significant correlations (p < .05, Holm-Bonferroni corrected for 26 comparisons), as did candidate auditory regions tgPCS, cIFS/G, aCO, FO, and cmSFG.

In a separate analysis, we inspected each individual subject’s task activation map of auditory WM (auditory 2-back) contrasted with visual WM (visual 2-back) to assess the robustness of sensory-biased WM activation in each candidate structure. Figure 4A shows maps for two representative subjects; the complete set of subjects is shown in Figure S2. Each region from Figure 3 was scored by two authors as *Strong* (1.0), *Weak* (0.5), or *Absent* (0.0) in each subject; Table S1 reports the average scores for each rater and region.

All regions with significant split-half reliability of vertex-by-vertex activation also were scored 0.7 or higher by both raters. Regions with significant split-half reliability and high mean ratings were thus identified as reliable sensory-biased WM regions. By *sensory-biased*, we mean that these regions have significantly stronger activation during visual WM than during auditory WM, or vice versa. Candidate visual-biased region bilateral preSMA, was low in split-half reliability (LH *r* = 0.55, corrected *p* = .112; RH *r* = 0.29, corrected *p* > .99) and moderate in visual scoring (LH Rater 1 score = 0.60, Rater 2 score = 0.53; RH Rater 1 score = 0.77, Rater 2 score = 0.63) and therefore there was not sufficient evidence to support classifying this as a sensory-biased region. All other regions with low split-half reliability were visually scored 0.30 or lower (Table S1) and were rejected as sensory-biased regions.

This resulted in a set of eight bilateral sensory-biased regions for subsequent analysis. Previously, we reported two bilateral visual-biased structures lying in the superior precentral sulcus (sPCS) and inferior precentral sulcus (iPCS) (Michalka et al., 2015; Noyce et al., 2017; Tobyne et al., 2017); in addition, a third visual-biased structure reliably occurs ventral and anterior to iPCS, lying in the middle inferior frontal sulcus (midIFS). We also observe bilateral auditory-biased structures lying on the transverse gyrus bridging precentral sulcus (tgPCS) and in the caudal inferior frontal sulcus/gyrus (cIFS/G), as previously reported (Michalka et al., 2015; Noyce et al., 2017). A third auditory-biased structure lies on the anterior central operculum (aCO) and a fourth occurs on the frontal operculum (FO). On the medial surface, an auditory-biased structure occurs in the caudomedial portion of the superior frontal gyrus (cmSFG). Table 1 summarizes the modality preference, location, and size of these regions. Each ROI was evident in at least fourteen of the fifteen subjects.

**Table 1:**
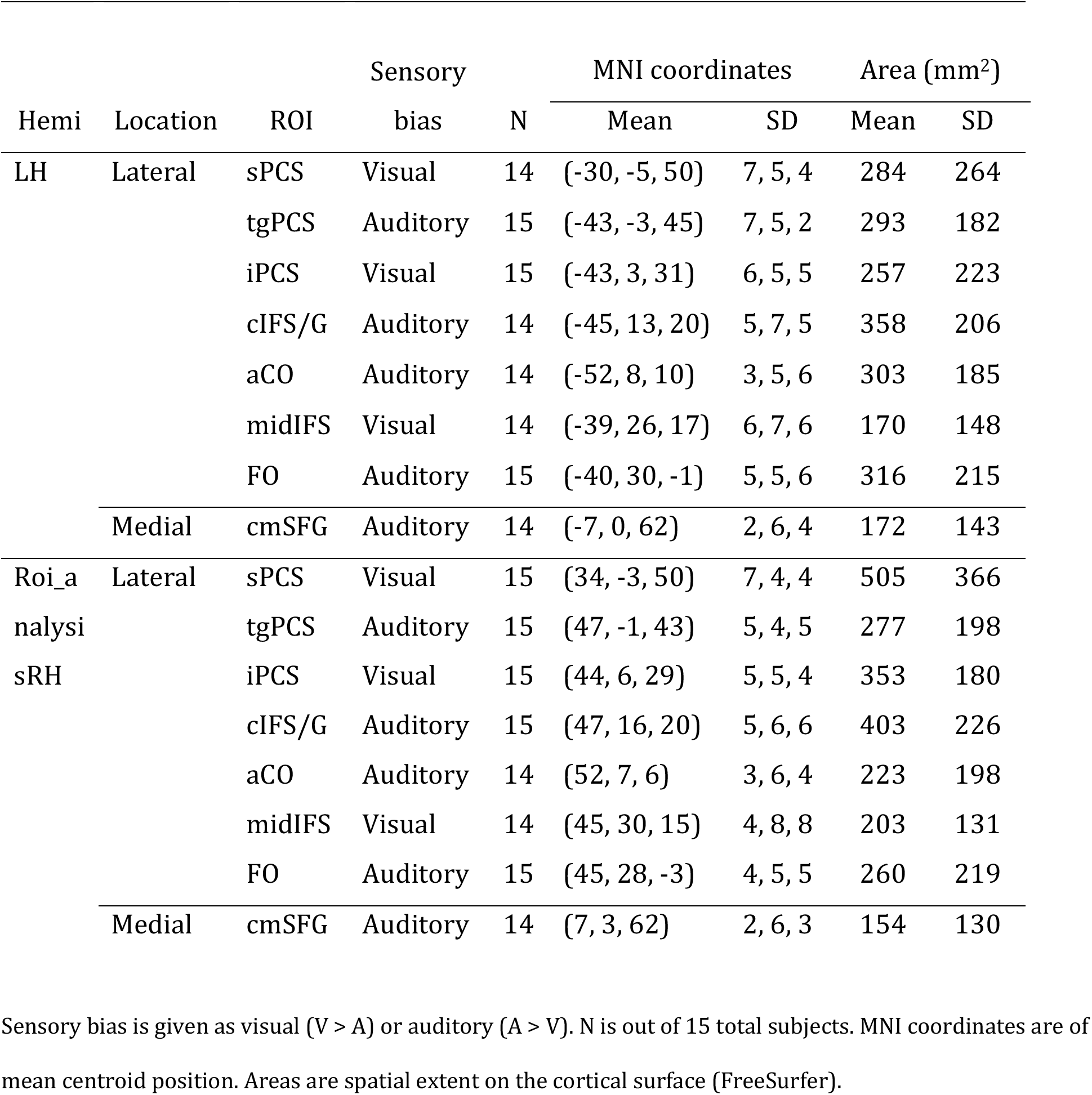
Size and position of sensory-biased ROIs

Each subject’s labels for the eight bilateral structures were projected into *fsaverage* space to create a probabilistic ROI map that visualizes anatomical consistency across subjects. Figures 4B and 4C show the degree of overlap of these labels for visual-biased and auditory-biased regions. For each structure, there is variability in its exact positioning across participants; moreover, this variability tends to be greater for more-anterior ROIs (particularly midIFS). Outlines in Figures 4B and 4C show the range of vertices that occur in at least 5 subjects (33%). Note that despite substantial positional variability in right midIFS (Figure 4B), that region was robustly identified in individual subjects (Table 1, Table S1, Figure S2).

### Quantifying sensory and WM drive in each region

For each sensory-biased frontal region, we computed the average % BOLD signal change in two contrasts: auditory vs. visual sensorimotor control, and auditory vs. visual 2-back WM (Figure *5*B, 5C). The contrast of sensorimotor control conditions captures the degree of sensory preference that each region exhibits in the absence of the 2-back WM task. We observe that the majority of regions (bilateral sPCS, iPCS, tgPCS, cIFS/G, aCO, and cmSFG, and right midIFS and FO) show a significant sensory preference. (All regions exhibit sensory drive in the expected direction, and all are significant before correction for multiple comparisons). However, in the presence of a working memory task, the degree of sensory specialization increases in most regions. Bilateral sPCS, iPCS, midIFS, tgPCS, cIFS/G, and FO, and and left aCO and cmSFG are all driven significantly more strongly by the 2-back task than by sensorimotor control. (Again, all comparisons are significant before correction for multiple comparisons.) We did not explicitly test the activation against zero in the 2-back condition as that would be “double dipping”; that contrast was used to define the regions under analysis.

**Figure 5.**
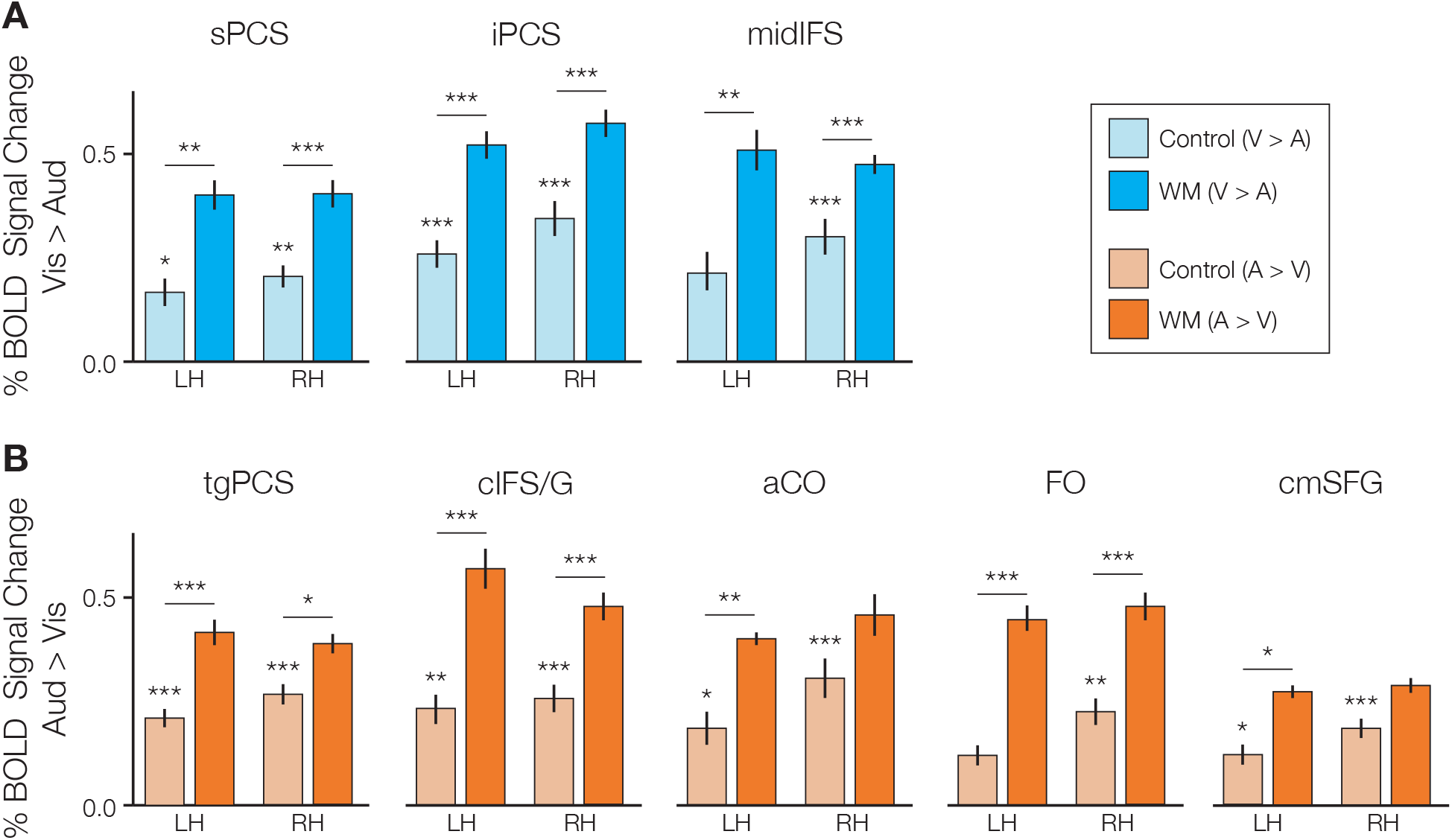
(A) Average sensory (light blue; visual sensorimotor control > auditory sensorimotor control) and WM (blue; visual 2-back > auditory 2-back) activation in visual-biased regions. Bilateral sPCS and iPCS and right midIFS are driven significantly in the absence of the 2-back task; bilateral sPCS, iPCS, and midIFS are driven significantly more strongly by the 2-back task than by the sensorimotor control. (B) Average sensory (light orange; auditory sensorimotor control > visual sensorimotor control) and WM (orange; auditory 2-back > visual 2-back) activation in auditory-biased regions. Bilateral tgPCS, cIFS/G, aCO, and cmSFG, and right FO are driven significantly in the absence of the 2-back task; bilateral tgPCS, cIFS/G, and FO, and left aCO and cmSFG are driven significantly more strongly by the 2-back task than by the sensorimotor control. *p < .05, **p < .01, ***p < .001; Holm-Bonferroni corrected for 16 comparisons.

### Network structure of visual-biased and auditory-biased regions

Above, we employed differential functional connectivity to identify candidate sensory-biased regions and then employed task data to test whether each candidate region was truly sensory-biased. Now, to test whether these confirmed sensory-biased regions, whose labels were drawn according to task activation boundaries, do indeed form discrete sensory-specific networks, we performed two resting-state functional connectivity analyses (n = 14, as one subject lacked resting-state data). First, we mapped each subjects’ individual sensory-biased frontal structures as well as broad posterior visual and auditory attention regions in order to define subject-specific ROI seeds. We then measured connectivity from each frontal structure (seeds) to the two posterior regions (targets). We ran a repeated-measures linear model with fixed effects of target region (pVis, pAud) and seed region sensory bias (visual, auditory), and random intercepts for subject and seed region. Connectivity to posterior regions varied strongly as an interaction between target region and seed sensory bias (F(1,13) = 248.8229, p < .0001), such that visual-biased regions were connected more strongly to pVis (r = 0.583) than to pAud (r = 0.440) and auditory-biased regions were connected more strongly to pAud (r = 0.604) than to pVis (r = 0.312). Overall connectivity was slightly stronger to pAud (r = 0.503) than to pVis (r = 0.415; F(1,13) = 30.736, p = .0002), and there was no main effect of seed region sensory bias (F(1,13) = 0.0006).

Second, hierarchical clustering analysis (HCA) was performed to more closely examine the network structure of these 2) ROIs. A distance matrix was constructed using each seed’s connectivity to each other region in the set (16 frontal and 4 posterior). HCA applied to these connectivity profiles confirmed that these regions assort into two discrete sensory-biased networks (Figure 6). All of the visual-biased regions (as identified by task activation) formed one network, while all of the auditory-biased regions formed the other network. A bootstrap test of the reliability of this behavior confirmed that on more than 97% of bootstrap samples, the auditory and visual subtrees were perfectly segregated from one another.

**Figure 6.**
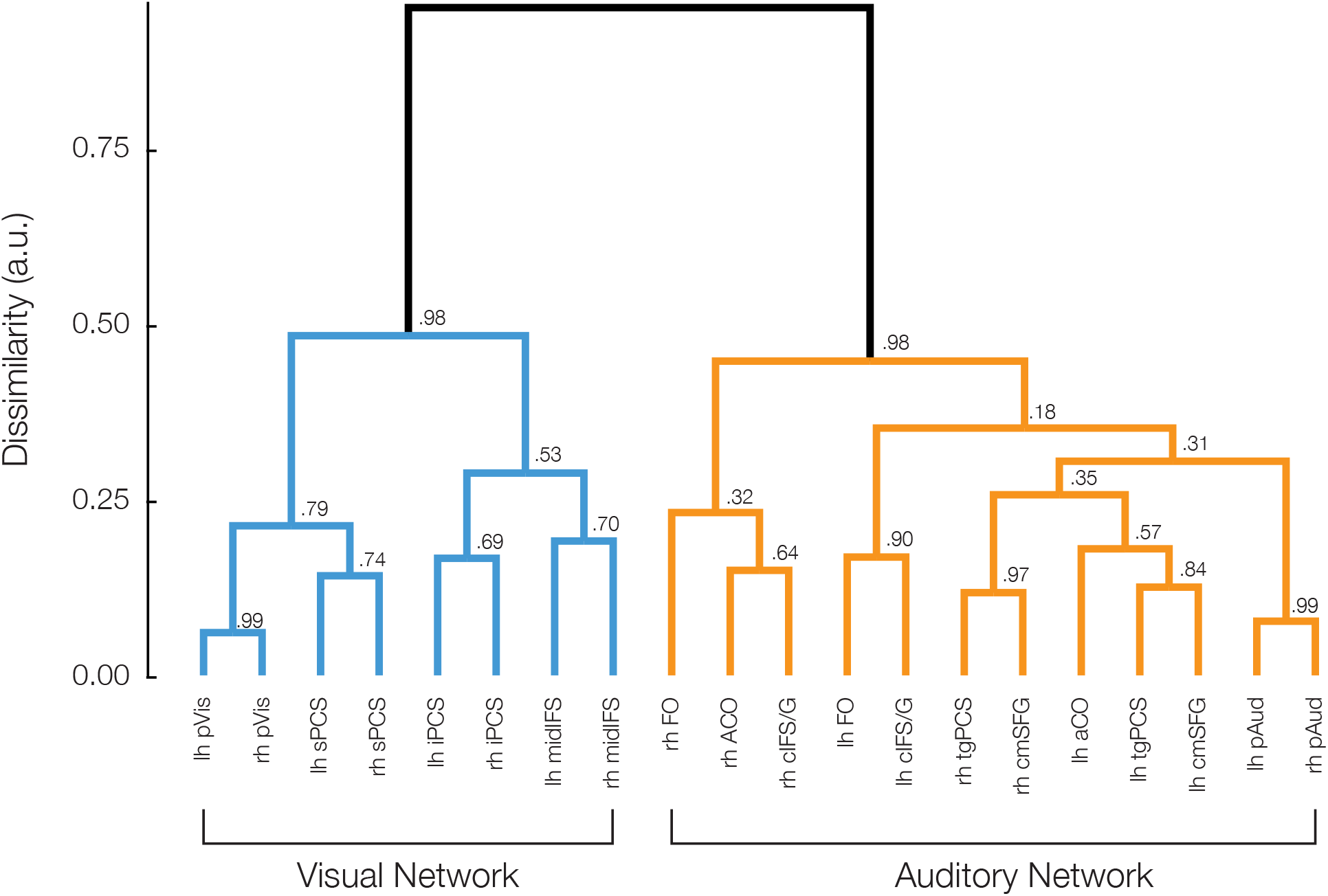
Hierarchical clustering analysis of seed-to-seed resting-state functional connectivity confirms that frontal visual and auditory WM structures assort into two discrete networks along with their respective posterior sensory attention structures. Values at each branch point show the reliability of each exact subtree, calculated as the proportion of bootstrapped datasets in which that subtree occurred.

Finally, we examined the WM-related behavior of regions that appeared in the differential functional connectivity analysis (Figure 3A), but that did not meet our criterion for reliable, sensory-biased working memory recruitment (Figure 3B, 4). These ‘rejected candidate’ regions included visual-connected candidate regions in bilateral aIns and preSMA, and auditory-connected candidate regions in rMFG and bilateral midIns, and dmCC. Each region was rejected because we failed to reliably identify sensory-biased task activation on the maps of individual subject hemispheres. In order to further characterize WM task activation in these regions, we drew ROI labels based on the group-average connectivity maps and projected them back into each individual subject’s anatomical space. We then computed the % BOLD signal change between 2-back working memory and passive exposure to the same stimuli for each region. Figure 7 shows the results.

**Figure 7.**
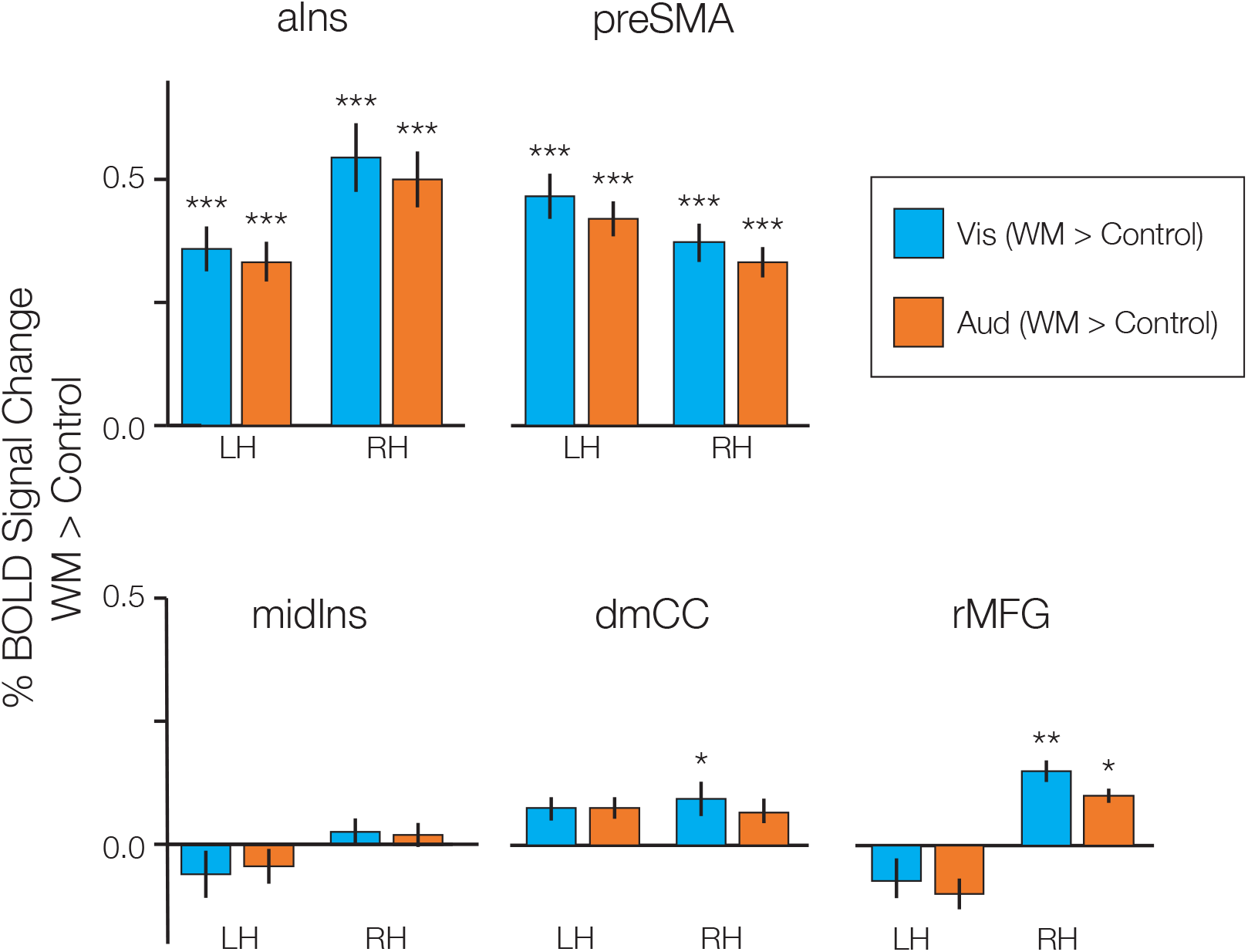
Task activation in rejected candidate regions; that is, extended network regions that do not show a sensory bias in WM recruitment. Rejected candidate ROIs are divided into members of the visual network (aIns and preSMA, top panel) and auditory network (midIns, dmCC, and rMFG, bottom panel). Bar plot shows mean BOLD % signal change in visual (blue) and auditory (orange) 2-back WM blocks, contrasted against sensorimotor control blocks. Note that aIns and preSMA, connected to the visual network, are robustly recruited in both WM tasks, while midIns, dmCC, and rMFG are at best minimally recruited in either. Error bars are repeated-measures standard error of the mean. * p < .05, ** p < .01, *** p < .001, Holm-Bonferroni corrected for 20 comparisons.

Bilaterally, aIns and preSMA, regions that are preferentially connected to the visual network, appear to be truly multisensory during WM, with robust (corrected *p* < .001) recruitment in both tasks. This is in sharp contrast to the regions that are preferentially connected to the auditory network (rMFG, midIns, and dmCC), which are not appreciably recruited in either WM modality. We ran a repeated-measures linear model with fixed effects of task (auditory WM; visual WM) and preferential connectivity (pAud > pVis; pVis > pAud), and random intercepts for subject, hemisphere, and region, and confirmed that task activation differed with preferential connectivity (F(1,13) = 81.3227, p < .0001) but not with either the WM task (F(1,13) = 3.1139) nor the task-by-connectivity interaction (F(1,13) = 0.6747).

## Discussion

Our results indicate that sensory modality is a driving factor in the functional organization of a substantial portion of frontal cortex. In both lateral and medial frontal cortex, we observed swaths of bilateral cortex, running caudodorsal to rostroventral, containing multiple regions that exhibit a preference for one sensory modality during N-back WM tasks. Our analysis began with thirteen bilateral candidate sensory-biased frontal lobe regions identified by intrinsic connectivity. Split-half reliability and scoring in individual subjects found that eight of these were sufficiently robust to draw individualized labels for further investigation. We tested their consistency of anatomical location, their degree of sensory and WM-specific drive, and their participation in sensory-specific resting state networks. In each individual subject, the frontal lobes in both hemispheres exhibited multiple regions that were activated more strongly for visual WM than for auditory WM and multiple regions that exhibited the opposite preference. Across our subjects, eight bilateral sensory-biased regions in frontal cortex exhibited a high degree of anatomical consistency, both in terms of stereotactic coordinates and in terms of relational position to other sensory-biased regions within the individual hemisphere (Figure 4B,C, Table 1). Although visual-biased and auditory-biased regions interleave across frontal cortex, hierarchical clustering analysis of resting-state data demonstrates that they form two distinct functional connectivity networks, one containing all of the visual-biased regions and the other containing all of the auditory-biased regions (Figure 7). In prior work, we had identified four bilateral sensory-biased frontal lobe regions, visual-biased sPCS and iPCS and auditory-biased tgPCS and cIFS/G (Michalka et al., 2015; Noyce et al., 2017). Here, we extend those findings to report four additional sensory-biased frontal regions, bilaterally. One new visual-biased frontal region was identified, midIFS, a region midway along the inferior frontal sulcus. Visual-biased preSMA, a region on the medial surface, exhibited some properties of the other visual-biased regions, but fell short of our criteria for a reliable sensory-biased region. Three new auditory-biased frontal regions were identified, aCO, on the anterior portion of the central operculum, FO, a region on the frontal operculum, and cmSFG, a medial surface region lying in the caudomedial portion of the superior frontal gyrus. On the lateral surface, visual-biased sPCS is the most caudal and dorsal region. Running rostroventral from sPCS are tgPCS (aud), iPCS (vis), cIFS/G (aud), midIFS (vis), and FO (aud). aCO lies caudoventral to cIFS/G. On the medial surface we reliably observed auditory-biased cmSFG. These results confirm the existence of these regions, and their sensory bias, as posited in Tobyne et al. (2017).

Although sensory modality is widely accepted to be a major organizing principle for occipital, temporal, and parietal lobes, only a handful of human or non-human primate studies have examined sensory modality as a factor in the organization of the frontal lobes (Barbas & Mesulam, 1981; Braga et al., 2013, 2017; Mayer et al., 2016; Michalka et al., 2015; Noyce et al., 2017; Petrides & Pandya, 1999; Romanski, 2007; Romanski & Goldman-Rakic, 2002; Tobyne et al., 2017). In human neuroimaging studies, activation within the frontal lobes is typically much weaker than activation within other cortical lobes. As a result, it is very common for frontal lobe studies to report only group averaged activity in order to increase statistical power. However, frontal cortex displays a high degree of inter-subject variability (Mueller et al., 2013; Tobyne et al., 2018), which may mask fine-grained sensory-specific organization. Numerous studies have concluded that frontal lobe organization is independent of sensory modality (e.g., Duncan, 2010; Duncan & Owen, 2000; Ivanoff et al., 2009; Krumbholz et al., 2009). Other studies have identified patterns of cortical organization based on biases towards visual versus auditory processing (Crottaz-Herbette et al., 2004; Fedorenko et al., 2013; Mayer et al., 2016) or connectivity (e.g. Blank, Kanwisher, & Fedorenko, 2014; Glasser et al., 2016; Braga, Hellyer, Wise, & Leech, 2017); however, none of these approaches have identified the pattern of interdigitated visual- and auditory-biased structures that is so striking in individual subjects here. More recent work has localized multiple demand regions within lateral frontal cortex with a higher degree of precision and less inter-subject smearing (Assem et al., 2020).

Our present analysis was initially guided by contrasting the intrinsic functional connectivity of the two visual-biased regions that we had previously identified, sPCS and iPCS, with that of the two auditory-biased regions that we had previously identified, tgPCS and cIFS/G. This analysis revealed not only the expected sensory-biased regions in temporal, occipital and parietal cortex and the four frontal seed regions, but also six regions on the lateral surface and insula and three structures on the medial surface. Examination of task activation led us to reject five of these nine candidate regions, which did not show consistent patterns of activity across subjects, while supporting the four others. Of these four regions, the three lateral structures, midIFS (vis), FO (aud), and aCO (aud) were each robustly observed in 80% or more of individual hemispheres. Our observed visual-biased and auditory-biased regions, particularly in caudolateral frontal cortex, tend to lie posterior and adjacent to the multiple demand regions found by Assem et al. (2020).

We tested the degree of pure sensory drive and of sensory-specific WM activation in each region of interest. The vast majority of regions exhibited both a significant preference for auditory or visual activity during sensorimotor control blocks, and a significant increase in activity during 2-back WM. These regions thus appear to be recruited both for perceptual processing, and also to support demanding cognitive tasks such as WM in their preferred modality.

One interesting result in the hierarchical clustering analysis is that within the visual network, the lowest-level clusters first tend to group the same regions in the two hemispheres, such as left and right sPCS, or left and right midIFS; in contrast, within the auditory network, the left cIFS/G and FO group together, as do the right cIFS/G, aCO, and FO; similarly, the left aCO, tgPCS, and SMA group together, as do the right tgPCS and SMA. This may be evidence for greater hemispheric specialization within the auditory network, perhaps related to language and/or speech processing, than in the visual network.

The contrast of resting-state functional connectivity from visual-biased and auditory-biased frontal seed regions generated some candidate regions (aIns, preSMA, midIns, rMFG, dmCC) that the task analysis rejected. This result indicates that the seed regions (sPCS, iPCS, tgPCS, cIFS/G) not only belong to visual-biased and auditory-biased attention & WM networks, respectively, but also to other functional networks. Note that univariate analyses may spuriously reject some regions that do discriminate among conditions (e.g. Harrison & Tong, 2009); however, the stimuli and task used here are not sufficiently well controlled to be good candidates for a multivariate classifier-based approach. Future work should more carefully assess this question.

Previously, we noted that the visual-biased frontal regions (sPCS, iPCS) exhibited a significantly greater responsiveness to the non-preferred modality than did the auditory-biased frontal regions (tgPCS, cIFS/G) (Noyce et al., 2017); that is, the visual-biased regions tended to exhibit a degree of ‘multiple demand’ functionality. In that work, we also characterized aIns and preSMA as full ‘multiple demand’ regions that were strongly recruited for WM tasks in both sensory modalities (see also Assem et al., 2020; Duncan, 2010; Duncan & Owen, 2000; Fedorenko et al., 2013). Here, we confirmed that aIns and preSMA respond robustly to WM tasks in both sensory modalities in individual subjects and thus reaffirm the claim of multiple-demand functionality. However, aIns and preSMA exhibited stronger resting-state connectivity with the visual-biased than with the auditory-biased frontal regions.

In contrast, the rejected auditory candidate regions, midIns, dmCC, and rMFG, exhibited little response to the WM demands in either sensory modality. Noyce et al. (2017) observed that the visual-biased frontal structures exhibited some degree of multisensory WM recruitment, while the auditory-biased frontal structures were strictly selective for auditory tasks. Our observation that regions with preferential connectivity to those structures show related asymmetries between the visual and auditory networks is consistent with that result. It may be that visual-biased frontal structures are more multisensory exactly because they are more strongly connected to modality-general WM regions.

The present data do not provide a basis for drawing any further conclusions about the functionality of the auditory-connected regions; however, it is worth noting that midIns and dmCC appear to anatomically correspond to regions identified as part of the speech production network (Bohland & Guenther, 2006; Guenther, 2016). Additionally, auditory-biased attention regions aCO, FO, and cmSFG, which were revealed here with a non-speech task, also appear to anatomically correspond to regions recruited when subjects are asked to listen to and repeat back spoken syllables (Bohland & Guenther, 2006; Brown et al., 2005; Guenther, 2016; Markiewicz & Bohland, 2016; Scott & Perrachione, 2019; Turkeltaub et al., 2002). The possible overlap of speech production regions and auditory-biased attention regions should be examined in individual subjects; however, given the auditory WM demands of the speech production tasks, such an overlap would not be surprising.

Taken together, these observations suggest that the visual-biased attentional network hierarchically merges into or at least overlays with a multiple-demand network, with the degree of visual-bias varying across the network, finally disappearing in aIns and preSMA. Similarly, the auditory-biased attentional network may overlay with substantial portions of the speech production network; however, further direct studies are required to confirm this. Further studies are also needed to better understand the specific functional roles contributed by each specific region in these extensive sensory-biased attention networks.

We used a group-average analysis of differential functional connectivity to guide ROI definitions from fMRI task-activation ROI definition in individual subjects. We find that an extended frontal network of eight bilateral sensory-biased regions is robust and replicable across individual subjects, and that both task activation and resting-state functional connectivity affirm the sensory-biased identities of these regions. We further find that these regions exhibit both sensory drive in their preferred modality as well as significant increases in activation during WM. These results highlight the importance of understanding cortical organization on the individual subject level, because this fine-scale structure can vary slightly across individuals, smearing or blurring group-level effects. Finally, our results provide support for an emerging hypothesis that the auditory and visual cortical processing networks support fundamentally different kinds of computations for human cognition, with the auditory network providing very specialized processing and the visual network participating more generally across a range of tasks.

## Acknowledgments

We thank Ilona Bloem, Allen Chang, Kathryn Devaney, Emily Levin, David Osher, and Maya Rosen for assistance with data collection, and David Beeler, Ryan Marshall, Ningcong Tong, and Vaibhav Tripathi for discussions and feedback.

