## Supplemental Material for "Extended frontal networks for visual and auditory working memory"

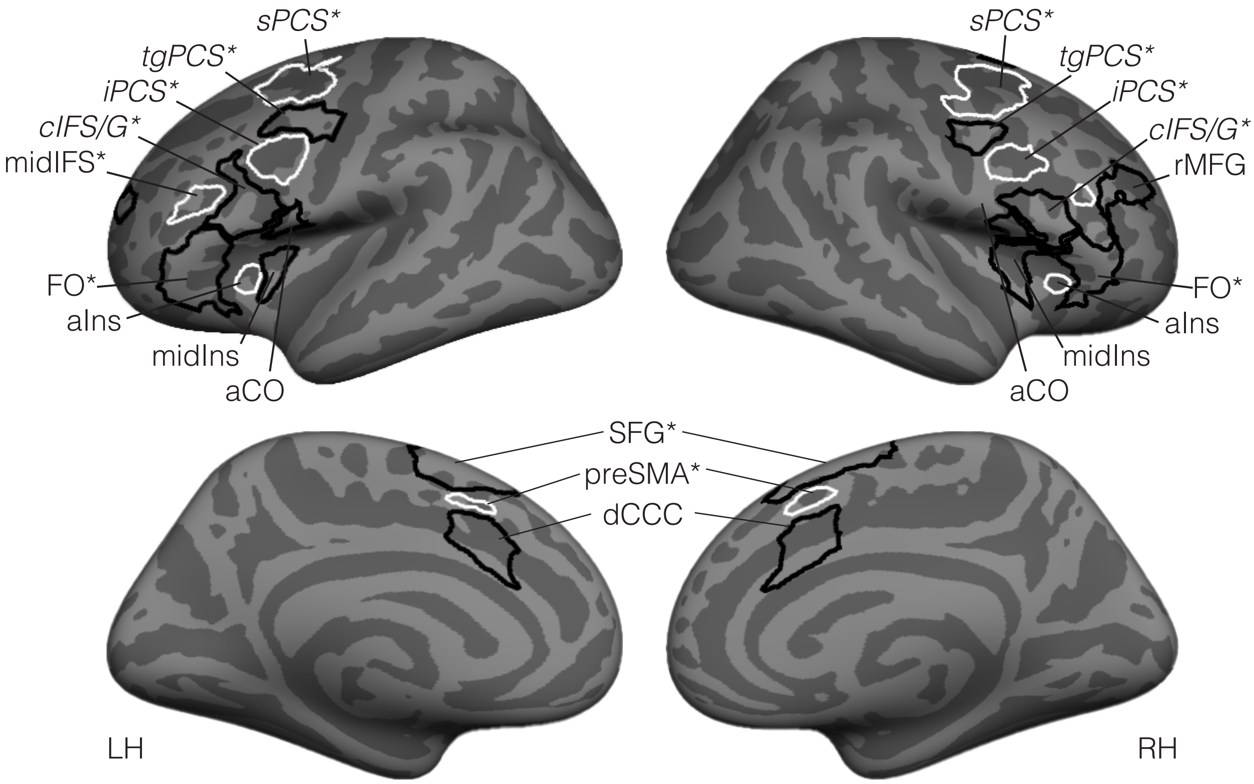


**Supplemental Figure 1.** Search spaces used for the split-half reliability analysis of candidate regions. Search spaces were drawn in the fsaverage template space based on differential connectivity (Figure 3). Spaces for candidate visual-biased regions are outlined in white; spaces for candidate auditory-biased regions are outlined in black.

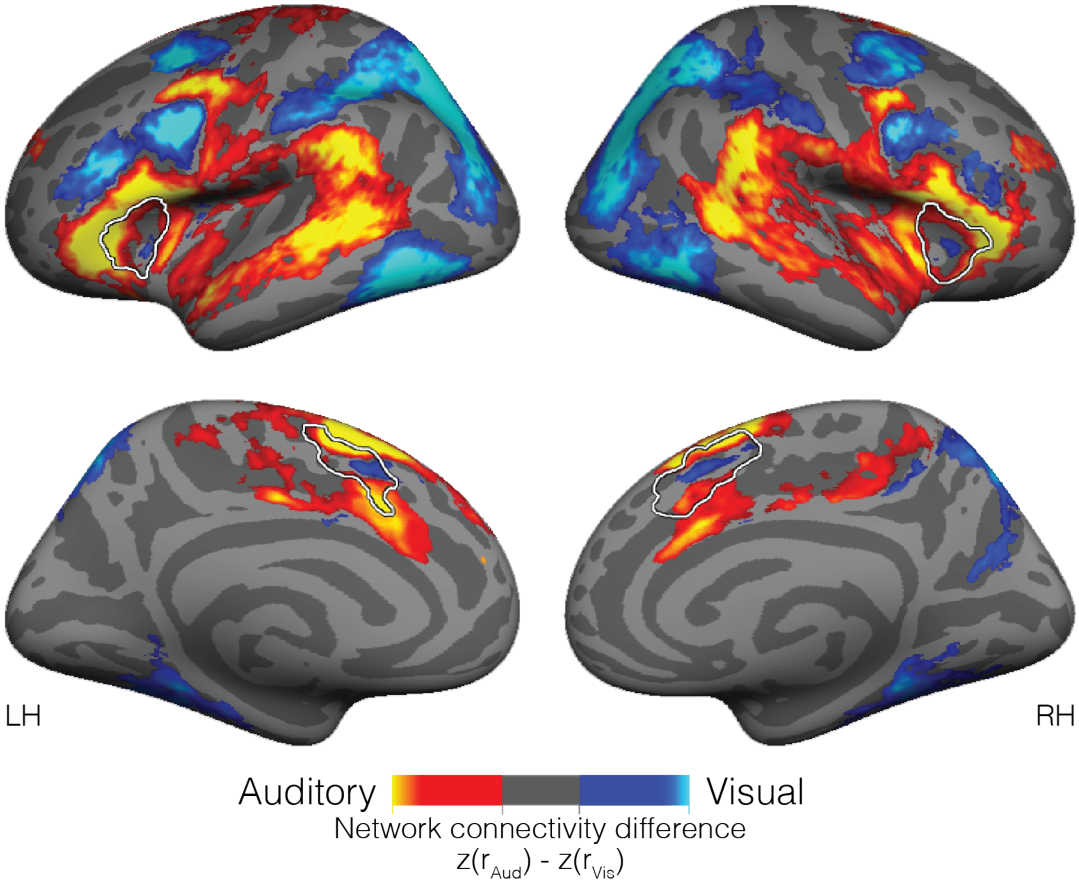


**Supplemental Figure 2:** Differential connectivity from frontal seeds (as in main Figure 1), with overlain outlines of the anterior insula and dorsal central cingulate cortex areas described in Noyce et al. (2017) as “multiple demand” regions.

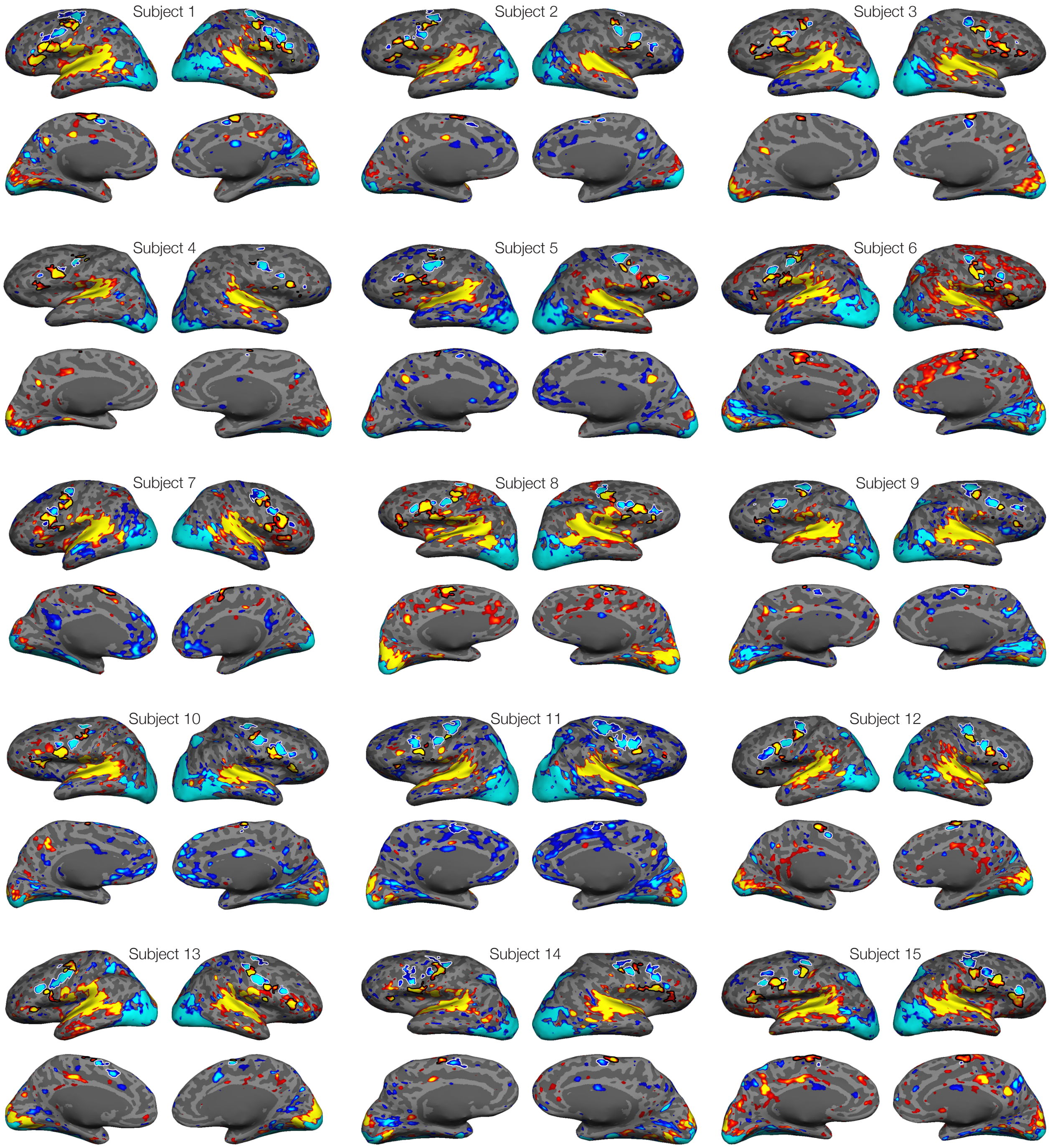


**Supplemental Figure 3:** Statistical maps of the contrast between auditory 2-back (hot colors) and visual 2-back (cool colors) in each of the fifteen subjects who participated in this study. Maps are thresholded at $p < .05$ (uncorrected). Each subject's frontal ROIs for subsequent analyses are outlined.

Table S1: Strength scores of candidate ROIs.

| Hemi | Location | Sensory bias | Candidate region | Rater 1 mean score (SD) | Rater 2 mean score (SD) |
|---|---|---|---|---|---|
| LH | Lateral | Visual | sPCS | 0.87 (0.30) | 0.77 (0.37) |
| | | | iPCS | 1.00 (0.00) | 1.00 (0.00) |
| | | | midIFS | 0.87 (0.30) | 0.83 (0.24) |
| | | | aIns | 0.50 (0.38) | 0.27 (0.26) |
| | | Auditory | tgPCS | 1.00 (0.00) | 0.97 (0.13) |
| | | | cIFS/G | 0.90 (0.28) | 1.00 (0.00) |
| | | | aCO | 0.90 (0.28) | 0.83 (0.24) |
| | | | rMFG | 0.23 (0.32) | 0.20 (0.25) |
| | | | FO | 1.00 (0.00) | 1.00 (0.00) |
| | Medial | Visual | preSMA | 0.60 (0.43) | 0.53 (0.44) |
| | | Auditory | cmSFG | 0.77 (0.32) | 0.73 (0.37) |
| | | | dmCC | 0.30 (0.46) | 0.13 (0.23) |
| RH | Lateral | Visual | sPCS | 1.00 (0.00) | 0.93 (0.18) |
| | | | iPCS | 1.00 (0.00) | 1.00 (0.00) |
| | | | midIFS | 0.93 (0.18) | 0.87 (0.23) |
| | | | aIns | 0.50 (0.38) | 0.27 (0.37) |
| | | Auditory | tgPCS | 1.00 (0.00) | 0.97 (0.13) |
| | | | cIFS/G | 1.00 (0.00) | 0.97 (0.13) |
| | | | aCO | 0.83 ( 0.31) | 0.73 (0.32) |
| | | | rMFG | 0.27 (0.37) | 0.20 (0.25) |
| | | | FO | 0.90 (0.21) | 0.87 (0.30) |
| | Medial | Visual | preSMA | 0.77 (0.43) | 0.63 (0.40) |
| | | Auditory | cmSFG | 0.80 (0.32) | 0.77 (0.32) |
| | | | dmCC | 0.27 (0.37) | 0.17 (0.31) |
